# Single-cell learning in *Stentor coeruleus* is governed by a fractional-order low-pass filter

**DOI:** 10.64898/2026.06.06.730631

**Authors:** Salvador Escobedo, Aline Moran, Frank Wu, Elvira Magaña, Akira Bowler, Karina Rodriguez, Gurleen Kaur, Julianna Mululu, Katlyn Benton, Aya Alkabbani, Ximena Garcia Arceo, Wallace F. Marshall

## Abstract

Single cells display a range of complex behaviors normally associated with a nervous system, including basic forms of learning like habituation. The giant ciliate *Stentor coeruleus* habituates to mechanical stimuli and shows many of the hallmark features characteristic of habituation in animal cells. When *Stentor* cells are mechanically stimulated by a predator or other stimuli, an action potential fires and leads to calcium-dependent contraction. When the same cell is repeatedly stimulated, it becomes less likely to respond, thus showing habituation. While the molecular basis of habituation in *Stentor* is not yet known, it has been shown to involve CaMKII, which also plays a key role in learning in neurons. The presence of an action potential, the role of calcium in the response, and the involvement of CaMKII in habituation, all suggest a potential deep conservation of learning mechanisms between single-celled protists and the neurons of animals. A number of different models have been proposed to explain habituation in a single cell but existing data in *Stentor* are unable to clearly rule out any of these models or favor others. Here we report a frequency domain analysis of habituation in which we measure the response probability of *Stentor* cells to pulsatile stimuli delivered at a range of frequencies. We find that the Bode plot of the frequency response resembles a classic low pass filter, with a flat passband at low frequencies, a clear corner frequency, and a linear roll-off. However, unlike standard low pass filter, the roll-off occurs with a slope of -30dB/decade, thus showing a fractional-order behavior. None of the existing models for habituation in *Stentor*, at least in their current form, predict this form of the frequency response, leading us to look for other explanations. We tested, and ruled out, a model based on a refractory period associated with the re-extension of cells following contraction. Inspired by methods used in analog circuit design to approximate fractional order systems using conventional lumped devices, we developed a model in which a series of distinct molecular species, such as different multimeric complexes of CaMKII, acting in parallel to inhibit the response, produce a fractional-order effect. The fractional order behavior of habituation in *Stentor* resembles the fractional-order behavior of adaptation in neurons, further supporting the idea that neurons may employ similar mechanisms for learning as were already present in unicellular eukaryotes prior to the evolution of metazoa.

## Introduction

Single cells can display complex behaviors of a type normally associated with the presence of a nervous system (Binet 1897; Jennings 1906; Vertosick 2002; Bray 2011; Baluska and Levin 2016; Lyon 2021; Ros-Rocher and Brunet 2023; Larson 2023; Wan 2024). Examples include decision making, food selection, predator evasion, anticipation of future events, gait coordination, and finding shortest paths during navigation of complex environments (Golding 2011; Saigusa 2008; Larson 2022; Nakagaki 2000). Single cells can also display basic forms of learning (Jennings 1902; Tang and Marshall 2018; Dussutour 2021; Gershman 2021; Kukushkin 2024).

Somehow, all of these processes must be mediated by machinery that exists in a single cell, which could include biochemical signaling networks, genetic regulatory networks, membrane potentials, or dynamics of cellular structures. Networks of interacting biomolecules are capable of quite complex functions including learning (Fernando 2009; Gunawardena 2022; Trifonova 2025). Here we will focus on the most basic form of learning, namely habituation, and investigate how it takes place in a single cell by studying the process in the frequency domain.

Habituation is a form of non-associative learning traditionally defined as a gradual decrease in response to a stimulus upon repeated application of the same stimulus (Harris 1943). Habituation is ubiquitous among animals, including both vertebrates and invertebrates, and studies in a wide range of different animals have found that habituation is usually associated with a set of highly conserved additional features known as the “hallmarks of habituation” (Thompson and Spencer 1966; Rankin 2009). These include the tendency for stronger stimuli to cause less habituation, and for prolonged training to cause the animal to retain its habituated condition for a longer period of time after the stimulus is removed. One of the most widespread hallmarks of habituation, seen in virtually all animals in which it has been tested, from invertebrates to humans, is the tendency for habituation to happen more rapidly and to a greater extent when the stimuli are provided more frequently Thompson and Spencer 1966; Rankin 2009). With respect to terminology, we note that the terms habituation and adaptation both refer to a response decrement to a continued stimulus. In animals, a response decrement is called adaptation if it occurs at the sensory level, whereas it is called habituation if it occurs within the central nervous system (Staddon 1996; Kuenen 1981; Torre 1995). In the context of cell signaling, adaptation refers to restoring a set-point following a perturbation (Ferrell 2016), while habituation refers to a progressively decreasing response to repeated stimulation (Smart 2024).

Habituation, together with many of its hallmarks including the effect of stimulus frequency, has been clearly demonstrated in the unicellular organism *Stentor coeruleus* (Wood 1970a). *Stentor* (Tartar 1961; Marshall 2021) is a giant cone-shaped ciliate roughly 1 mm in length, that attaches to substrates like pond plants through a holdfast at one end, and creates a feeding current to capture prey by means of a ring of cilia at the other end, known as the membranellar band. When the *Stentor* cell receives a mechanical stimulus such as fluid flow or contact with a moving object, it can contract down into a roughly spherical shape, while retaining its attachment to the surface via its holdfast (**Figure 1A**). This contraction is thought to be a mechanism for predator evasion, since it pulls the bulk of the cell body away from its previous position and moves it down to the substrate. The entire process is extremely rapid, taking place in tens of milliseconds (Jones 1970; Newman 1972), and is driven by contraction of myonemes, cables made of the EF-hand calcium binding protein centrin (Huang and Pitelka 1973; Maloney 2005), which undergoes a conformational change that leads to a rapid shortening of filament length. The contraction is triggered by entry of calcium into the cell via voltage gated calcium channels (Wood 1982), whose activity is triggered by an action potential (Wood 1970b; Wood 1988). The action potential that triggers contraction depends on a membrane depolarization that occurs when the cell is touched (Wood 1970b), presumably generated by a channel-linked mechanoreceptor. The identity of the receptor is not yet known, although pharmacological evidence suggests it may be a member of the nicotinic acetylcholine receptor family (Wood 1977; Wood 1985; Rajan 2026).

**Figure 1.**
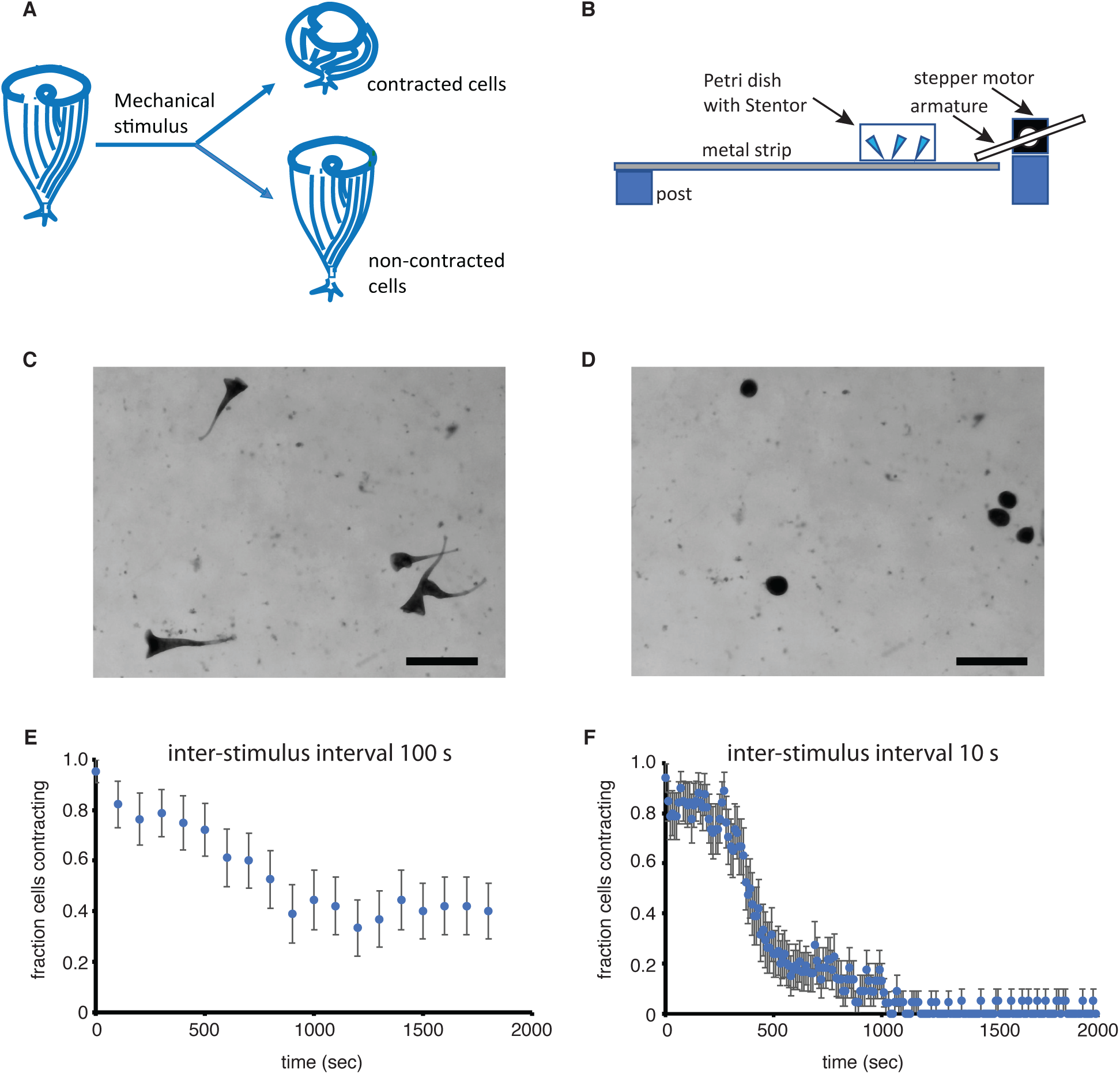
Habituation in *Stentor coeruleus*. **A.** Diagram of a *Stentor coeruleus* cell, with a ring of cilia at one end and a holdfast at the other, where it attaches to substrates like pond plants. When the cell is undisturbed it extends into a cone shape and feeds by creating a flow of fluid using the ring of cilia. When disturbed by a mechanical force, the cell can contract into a spherical shape to avoid predation. When disturbed repeatedly with the same level stimulus, the cell will gradually stop responding, thus manifesting habituation. **B.** Device used to measure habituation in *Stentor*. **C.** Image of cells pre-contraction. Scale bar 1 mm. **D.** Image of cells post-contraction. Scale bar 1mm. **E.** Fraction of cells contracting versus stimulus number, for stimuli presented once every 100 seconds. N= 22. **F.** Fraction of cells contracting versus stimulus number, for stimuli presented once every 10 seconds. N= 20.

When a population of *Stentor* cell is repeatedly exposed to the same mechanical stimulus, the probability of the cells contracting will gradually decrease (Wood 1970a). This response decrement does not simply represent exhaustion, because cells habituated to a weak stimulus will still contract in response to a strong stimulus (Wood 1970a) or to pulses of light (Wood 1970a). When the stimulus is stopped, the cells will gradually recover their ability to contract in response to future stimuli (Wood 1970a). In addition to showing the response decrement by which habituation is defined, this process in *Stentor* also displays many of the classical habituation hallmarks.

Habituation in *Stentor* happens more rapidly and to a greater extent with weaker stimuli or when stimuli are applied more frequently (Wood 1970a; Rajan 2023a).

The molecular pathways that implement habituation in *Stentor* are not yet understood, but recent proteomic, transcriptomic, pharmacological, and RNAi experiments have demonstrated that members of the calmodulin dependent protein kinase II (CaMKII) family of protein kinases are involved in the process (Rajan 2026). CaMKII also plays a central role in learning in animal neurons (Hell 2014; Lisman 2017; Bayer 2025), suggesting a deep conservation of cellular learning mechanisms.

Understanding how habituation occurs in *Stentor* could thus represent a way to understand the evolution of learning mechanisms in metazoa as well as giving insights into the ubiquity of basal cognitive processes across all of life (Lyon 2015; Wan 2021; Levin 2021; Reid 2023; Ros-Rocher 2023).

Several previous theoretical studies have produced models that can account for habituation in *Stentor*. Using a modeling framework drawn from signal processing, Smart et al (2024) showed that a model based on convolution of the input signal with an exponential decay kernel (essentially a one pole low pass filter) to produce a continuously varying output that serves as an internal variable whose magnitude determines the probability of the system responding to a pulsatile stimulus. This model was able to account for response decrement and many of the habituation hallmarks, but required multiple filter stages to be chained together in order to account for the full set of habituation hallmarks (Smart 2024). Gershman (2024) developed a model for habituation based on a Gaussian estimation process in which a filter is used to estimate the distribution of stimulus magnitudes that are likely to be produced in the environment which is then used as a basis for a policy of whether or not to respond to particular new stimulus. This simple modeling framework was highly effective in predicting many of the hallmarks of habituation, suggesting that habituation may indeed be a form of estimation process, in which the organism is attempting to estimate some feature of its environment that determines its policy for how to respond to future stimuli. The Gershman and Smart models are constructed at an abstract level and are agnostic about how they would be implemented with molecules. Eckert et al used a network enumeration strategy to identify signaling networks capable to demonstrating hallmarks of habituation (Eckert 2024). This model is closer to a molecular model since the nodes in the network represent signaling molecules. Finally, Rajan et al developed a biochemical model that combines a mechanosensitive receptor, an ion channel, a voltage gated calcium channel, and activation-dependent receptor inactivation and/or degradation (Rajan 2025).

All of these models were successful in explaining not only the habituation process itself but also many of the hallmarks of habituation. In particular, all four of these models make the basic prediction that when stimuli are provided at a higher frequency, meaning that less time elapses between successive stimuli, habituation is more effective and leads to a decreased response once the system has reached steady state. Here, we extend this analysis beyond just showing that habituation works better at higher frequency, and measure the exact form of the frequency dependence. This general strategy of measuring how a system responds to an input as a function of frequency is known as a frequency domain analysis, and has been applied to cell behaviors such as cell motility and neurite outgrowth (Ueda 1983; Odde 1995; Odde 1998; Uchida 1999; Silva 2006), but not, to our knowledge, to cell learning. Our approach is to view the cell like a low-pass filter in the sense that when inputs (stimuli) are delivered at sufficiently low frequency, they are transmitted to the output (contraction), such that every stimulus input gives a contraction output, but when frequency increases, it becomes more and more likely that a stimulus will not produce a contraction, which in the filter analogy means that less of the input signal is transmitted to the output. Within this framework, we can characterize the frequency dependence in terms of a gain versus frequency plot that represents the transfer function of the filter.

There are four reasons why we think it may be useful to characterize the behavior of a cell in terms of signal processing concepts such as filters. First, signal processing concepts have been extensively used to model the learning process, including habituation in single cells (Gunawardena 2022; Gershman 2024; Smart 2024). By measuring the frequency response of cells as they undergo habituation, it would become possible to obtain experimental data in a form that can be directly understood in terms of such abstract conceptual models. Second, as noted above, there are currently a number of models, presented at varying levels of biological detail, that can explain habituation in single cells. These models can, to varying extent, explain a number of known experimental results. Although all of these models share the prediction that higher frequency should lead to more extensive habituation, they differ in the exact dependence on frequency. By treating cells like filters and measuring their frequency response, it may be possible to discriminate among some of the competing models. Third, standard methods for representing filter transfer functions provide a compact way to represent a complex dataset using a small number of parameters. If such models can describe the response of cells to inputs of varying frequency, it would then be possible to represent the effect of various perturbations in terms of the small set of parameters of the filter transfer function. Finally, at a more philosophical level, one of the features of living matter compared to non-living natural objects is the importance of processing and storing information. By trying to understand cell learning in terms of signal processing, we open a new window into information transmission by living matter.

## Results

### Measuring the frequency response of Stentor habituation

In order to measure the frequency response of *Stentor*, we used a device we have previously described (Rajan 2023b), in which the *Stentor* cells are placed in a petri dish on a metal strip, which is then tapped by an aluminum armature attached to a stepper motor (**Figure 1B**). The motor is under computer control allowing the frequency and force of the stimuli to be varied. When they are not disturbed, *Stentor* cells take on an elongated cone shape (**Figure 1C**), but when tapped by the motor, they contract into a ball (**Figure 1D**). We then monitor the response of individual cells over time to assess habituation. **Figure 1E** shows the response of cells tapped once every 100 seconds (i.e. at a frequency of 0.01 Hz), showing that they gradually reduce their probability of contracting, from above 90% down to around 40%. This rate and level of habituation is consistent with what has been reported previous (Wood 1970a; Rajan 2023a; Rajan 2026). **Figure 1F** shows a different experiment in which cells are tapped once every 10 second (i.e. a frequency of 0.1 Hz). In this case, although the cells started out contracting with the same initial probability as in the 100 second experiment, habituation was more extensive, with the contraction probability eventually dropping below 10%.

Using this experimental paradigm, we can define the frequency response in terms of the final probability of contracting, once habituation is achieved, as a function of the frequency with which the dish is tapped by the device. Our goal is to use measurements of this type to define the filter characteristics of the cell, thus producing a new type of data to compare with theoretical models of habituation.

First, we will define how we are interpreting our experiment in terms of a filter. Low pass filters are classically described by a Bode plot (**Figure 2A**), which is a log-log plot that describes the gain of the filter versus the frequency of the input. To generate such a plot, one first calculates the gain at each frequency, defined as the magnitude of the output divided by the magnitude of the input, such that if the input and output magnitudes are Mi and Mo respectively, the gain is Mo/Mi. For our measurements, since we are applying a binary input and looking at a binary output, the gain is simply the fraction of input stimuli that generate a contraction at any given frequency of input, which we denote P_contract_(f). The gain at zero frequency, known as the DC gain, is defined in our case as the probability that a cell will contract upon the first stimulus if it has not been previously stimulated. Formally this would require us to isolate cells from a stimulus for an infinite period of time, but in our experiments we let the cells sit for a four hour rest period before beginning the experiment.

**Figure 2.**
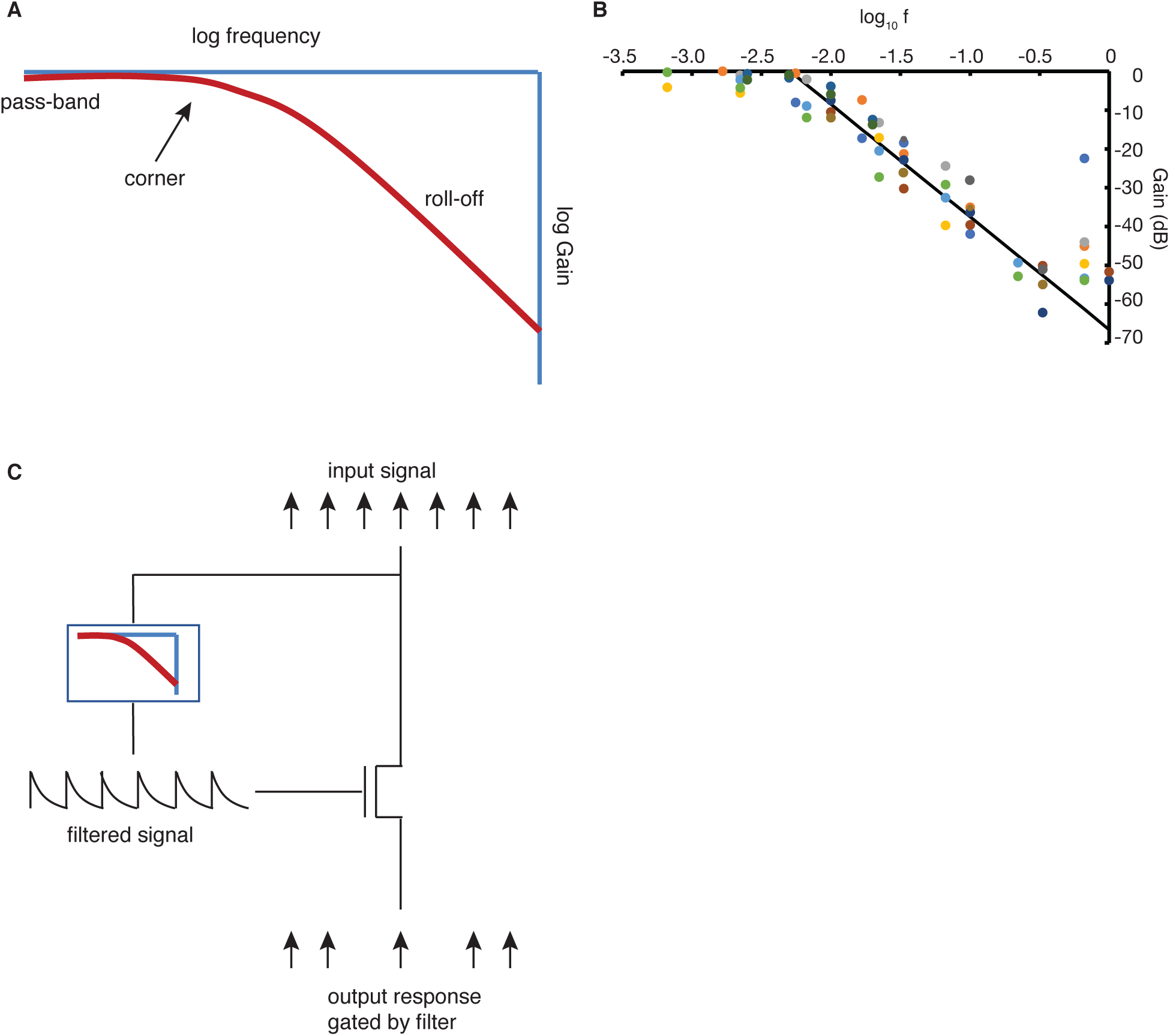
Frequency domain analysis of *Stentor* habituation. **A.** Diagram illustrating the general features of a low pass filter transfer function. Gain is expressed in unites of decibels, and frequency is plotted on a log scale. **B.** Experimentally measured transfer function for *Stentor*. Each marker corresponds to a habituation experiment like those of **Figure 1E,F**, in which the position of the marker corresponds to the probability of contraction once the cell has reached its final steady-state habituation level. Marker colors correspond to different batches of cells analyzed on different days. Gain is determined by dividing the steady state response probability by the average response probability at the start of each experiment in that batch, prior to habituation, which is taken as representing the DC gain. The gain in decibels is 20 times the log base 10 of the ratio of final to initial response probability. Line represent a best fit line to data in the frequency range 0.005-0.33 Hz, with slope 29 dB/decade and Y intercept 67 which corresponds to a corner frequency of 0.005 Hz. **C.** Interpreting the transfer function based on a model in which the cell responds to the stimulus with a linear filter that drives changes to an internal variable based on past stimuli, and which then modulates the cell response to incoming stimuli.

To generate the Bode gain plot, we normalize the gain at any given frequency P_contract_(f) by the DC gain P_contract_(0), and then express that normalized gain ratio in units of decibels (dB) according to the relation Gain=20*log10(P_contract_(f) / P_contract_(0)). The gain in dB is then plotted against the log10 of the frequency.

What makes a filter like that described in **Figure 2A** a low pass filter is that at sufficiently low frequencies, the input signal is propagated through the filter without attenuation, such that the gain is 0 dB. Once some threshold frequency, known as the corner frequency, is exceeded, the gain starts to decrease. To be more precise, the corner frequency is usually defined as the frequency at which the gain drops below -3 dB, correspond to an output whose amplitude is reduced by a factor of 2. Beyond the corner frequency, the gain typically drops linearly with a slope that depends on the details of the filter circuit. For the simplest type of low pass filter, known as a “one pole filter”, which is usually implemented by a single capacitor/resistor combination where charge on the capacitor decays as a single exponential, the gain decays with a slope of 20dB per decade. Sharper roll-offs can be achieved by chaining filters together to make a multi-pole filter, in which case the gain still decays linearly but with a slope that is an integer multiple of 20 dB/decade. Using this framework for describing the frequency domain response of a system, we set out to measure the frequency response of the *Stentor* cell in terms of its corner frequency and roll-off slope.

In order to measure the transfer function as per **Figure 2A**, we carried out a large number of separate habituation experiments, each like those shown in **Figure 1E** and **Figure 1F**, at a range of different stimulus frequencies. In each case, stimuli were applied using a stepper-motor based apparatus (Rajan 2023b) which allows us to control the periodicity of the mechanical stimulus. For each experiment, stimuli were applied at a constant frequency and taps are repeated until the response reaches a steady state (see Methods). Finally the last ten stimulus timepoints were analyzed and the average probability of response per cell per timepoint calculated. For higher frequencies at which the response probability becomes very small, we analyzed an increased number of timepoints to better estimate low-probability responses (see Methods for details). Once the final steady state response probability is determined for a given experiment, this response was normalized by dividing it by the average initial contraction probability, the ratio converted to dB, and then plotted as a single point on the gain-frequency plot.

Figure 2B shows the result of 64 separate habituation experiments plotted together on a single gain-frequency plot. Color of the markers indicates experiments done with different batches of cells on different days. In this plot, gain is calculated relative to the contraction probability at zero frequency, i.e., the case in which cells have not previously seen any stimulus. The resulting plot shows a clear low-pass filter behavior. At low frequencies, the gain remains close to 0 dB for frequencies below about 0.005Hz (i.e. once every 200 seconds), meaning that the response probability is the same as what we have defined as the response probability for zero frequency, namely, the response probability of naïve cells that had not previously been stimulated. This matches the qualitative observation previously reported that for sufficiently low frequency stimulation, habituation does not occur. As frequency is increased, the plot shows a clear corner, with the gain starting to decrease as the frequency reaches approximately 0.005 Hz, corresponding to stimuli provided at a period of ∼200 seconds.

As frequency is further increased beyond the corner frequency, the roll-off shows a visibly obvious linear decay. Fitting a straight line to all the data in the range 0.005-0.33 Hz, excluding the highest frequency data for which the number of observed contractions is extremely low making the response probability unreliable, gives a best-fit slope of -29 dB/decade. The linearity of this decay is notable because of its resemblance to a classical electronic low pass filter. However, the slope of ∼30 dB/decade is not something that would normally be seen with a standard low pass filter made from electronic components, which would either be -20 or -40 dB/decade, as will be discussed below. From the slope and intercept of the best fit line, we can calculate the intercept on the frequency axis, which is an indicator of the corner frequency, which comes out to be 0.005 Hz, consistent with the visual evaluation of the plot.

One potentially confounding element in this experiment, in terms of relating the results to standard signal processing theory, is that in our experiments, both the input and the output are sequences of impulses. The input is a sequence of discrete stimuli whose duration is short compared to the timescale of the observe, and hence can be treated like impulses. The output is a series of discrete contraction events that take place at times corresponding closely to the arrival times of input impulses. We can thus represent the input by a series of impulses (Dirac delta functions), known as a Dirac comb, while the output is a similar series of impulses, all of the same magnitude (given that contraction is an all or none event), in which some of the impulses may be missing. This is not what one would normally obtain if a Dirac comb is fed into a low pass filter. Instead, low pass filtering a Dirac comb should produce a continuously varying output function that represents a smoothed version of the comb, that is to say a series of peaks that are smoothed out along the time axis, with one peak per impulse in the input function. The shape of each peak is described by the impulse response of the filter, which for the simplest type of filter, known as a one-pole low pass filter, would be an exponentially decaying function whose time constant depends on the corner frequency of the filter. The reason that we don’t observe such a function is that we are not measuring output of the filter itself, but rather, a gated version of the input sequence, in which the probability that an input impulse produces an output impulse depends on the current output of the filter (see Figure 2C). This framework is highly similar to that of the Smart et al (2024) model in which a linear filter determines an internal variable that is then used to determine a response probability. Unlike the representation used by Smart et al (2024) in our case we equate the response probability with the filter output without invoking any additional intervening functions. In any case we are just using the filter analogy as a way to describe the observational data, without making a strong claim about the internal molecular circuitry that may be generating that data. We will address the latter point later.

### Does cell extension set the corner frequency?

The most fundamental characteristic of any low pass filter is its corner frequency since this determines the range of inputs that will be either passed or rejected by the filter. Typically the corner frequency is set as the inverse of the time required for some process that takes place inside the filter. For example in a simple electronic filter, the corner frequency is set by the time it takes to discharge a capacitor through a resistor. Measuring the corner frequency and thus the timescale can give clues about the internal processes taking place inside the filter.

Based on the data in Figure 2B, the corner frequency of 0.005 Hz, implies a timescale of approximately 200 seconds. Among known cellular processes related to contractile behavior that take place at such a timescale, an obvious candidate is the re-extension of the cells after contraction. While contraction itself occurs on the ten millisecond timescale (Newman 1972), driven by conformation changes of centrin making up the myonemes, cell re-extension is much slower, taking more than a minute, driven by sliding motion of the cortical microtubules (Huang and Pitelka 1973). In past experiments to study habituation in *Stentor*, the stimuli were usually applied at most once per minute (Wood 1970; Rajan 2023a), due to the concern that cells might not be able to respond to stimuli that occurred while the cell was still in the process of re-extension. In order to get a better idea of the actual timescale of re-extension, we imaged individual cells after a mechanically induced contraction and plotted their length versus time in Figure 3A. Although there is substantial variability between cells, it is clear that most cells reach their final length in a little over 100 seconds, with a half time of around 60 seconds, confirming prior assessment that re-extension occurs on a timescale of a minute.

**Figure 3.**
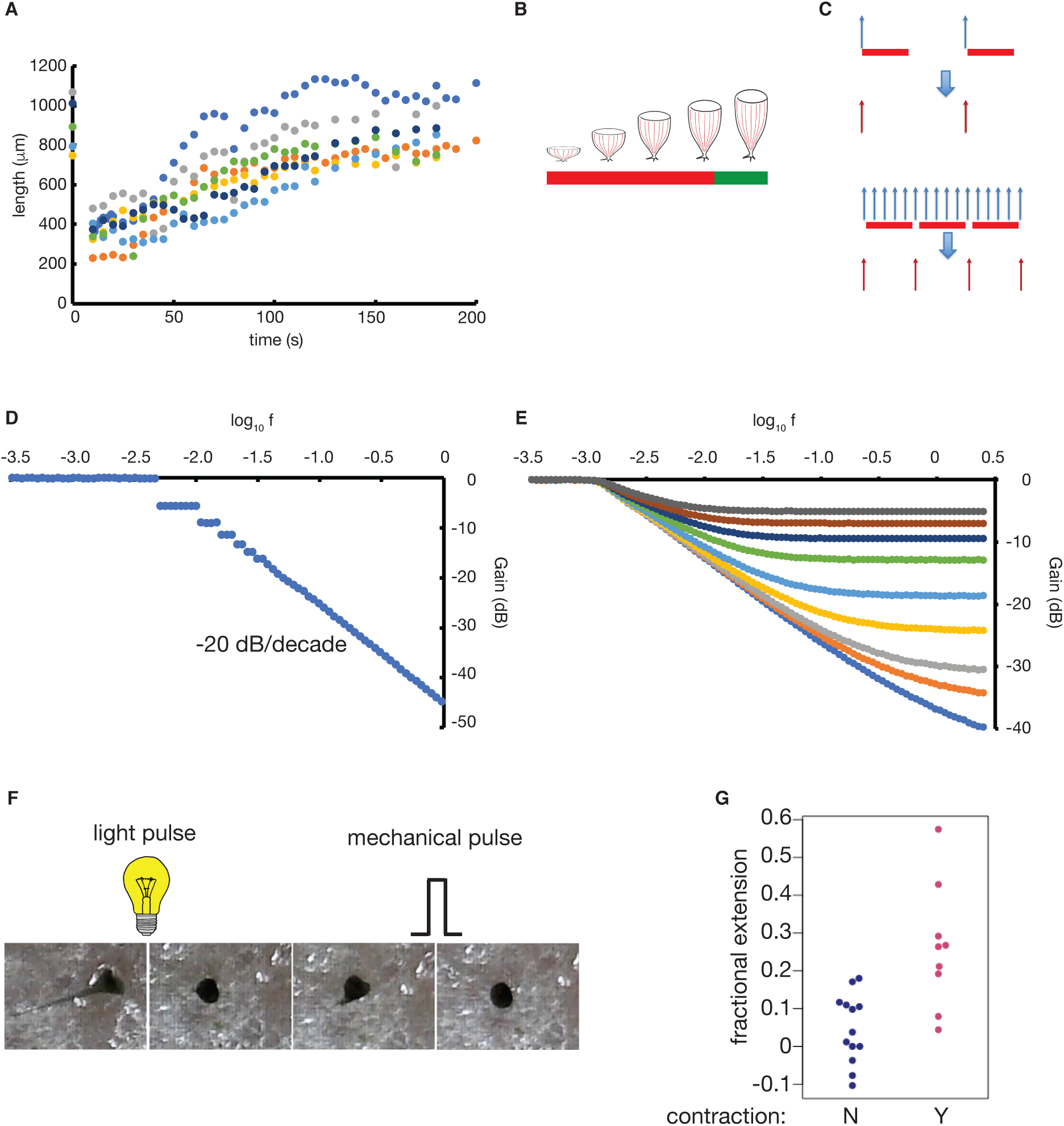
Testing a model for low pass filtering based on cell extension-dependent refractory period. **A.** Length versus time during re-extension of individual *Stentor* cells following a mechanical stimulus that caused them to contract. **B.** Diagram of cell extending showing a potential refractory period indicated by a red bar, followed by return to the responsive state as indicated by the green bar.. **C.** Diagram of a model with an absolute refractory period. Vertical arrows indicate individua stimuli and their associated responses. After responding to a stimulus, the cell enters a refractory state indicatd by the red bars. Only when the time period of the refractory period has elapsed can new stimuli trigger a response. **D.** Simulated filter transfer function for a probability of contraction in the responsive state of 0.9 and a probability of contraction in the refractory state of 0. **E.** Model prediction in which the refractory period was allowed to vary slightly between iterations and in which there was a non-zero probability of contraction while in refractory period. Simulations were performed with an average refractory period of 500 s and a standard deviation of 200 s. Curves show results for difference values of the probability of response during the refractory period, in order starting from the lower curve 0, 0.01, 0.02, 0.05, 0.1, 0.2, 0.3, 0.4, 0.5. **F.** Strategy for testing for a refractory period. Cells are first induced to contract in response to a pulse of intense light, then allowed to begin re-extending, and then subject to a mechanical stimulus at varying times during re-extension to determine when the cells become able to contract. **G.** Fractional extension denotes the fraction of total cell length that was re-established by the time of the mechanical stimulus. Contraction denotes whether cells did or did not contract when the mechanical stimulus was applied. Only cells that were extended prior to the light pulse and contracted after the light pulse were included in this analysis.

We consider the possibility that cells might enter a refractory state upon contraction and not be able to respond to stimuli until they had re-extended past some minimal length (Figure 3B). This could be the case if, for example, the centrin-based myonemes needed to be physically stretched in order to reset the conformational state of the centrin molecules back to one capable of mediating a contraction. What would the ramifications be of such a refractory state for the frequency response? As illustrated in Figure 3C, if stimuli were to arrive at a sufficiently low frequency, the refractory period would have no effect, because the cells would have fully re-extended before the next stimulus arrives. If, on the other hand, stimuli were arriving at a sufficiently high frequency, then some stimuli would end up being ignored because they would arrive while the cell was still in the process of re-extending, and thus in a refractory state.

Re-extension would thus lead to a low-pass filtering effect. As derived in Methods, in the simplest case in which cells are completely non-responsive for some fixed refractory period after contraction, set by the re-extension time, the frequency response would be given by

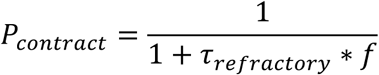

Where f is the frequency of the stimulus and 1_refractory_ is the time after contraction during which the cell is non-responsive (as diagrammed by the red bars in Figure 3B**,C**). This equation is exactly the equation of a one-pole low-pass filter with a corner frequency of 1/1_refractory_. The slope of the roll-off is thus predicted to be 20 dB/decade. This equation strictly holds only when a stimulus occurring outside the refractory period induces a response with probability 1. However, simulations with other response probabilities also show the same general behavior, with a corner frequency at 1/1_refractory_ and a linear roll-off with slope 20 dB/decade (see for example Figure 3D). If we relax the assumption that the cell cannot respond at all during the refractory period, we can obtain a series of transfer functions shown in Figure 3E. In this case, the frequency response drops after the initial corner frequency, but then eventually reaches a plateau at some lower response probability equal to the response probability during the refractory period. In between, the response falls off with a curve whose slope is less than 20 dB/decade. It is thus not possible, in this model, to obtain a roll-off of 30 dB/decade.

In order to explain a corner frequency of 0.005 Hz the refractory period would need to be roughly 200 seconds. Based on Figure 3A, this is the time scale at which cells have just begun to reach their full extension, and thus in order for the model to hold, cells would have to be non-responsive until they are essentially fully re-extended.

In order to test whether cells are indeed transiently non-responsive during mos or all of their re-extension period, we applied stimuli to cells that were still in the process of re-extending, and then asked how the response probability might change as a function of the degree of re-extension. Using a mechanical stimulus to induce contraction, and then using a second mechanical stimulus to probe the response during re-extension, is potentially problematic due to the fact that cells can start to habituate even after just one stimulus (Wood 1970a; Rajan 2023a). However, we can take advantage of the fact that while cells can also be induced to contract in response to bright pulses of light (Wood 1973), the two stimuli do not cross-habituate (Wood 1973), meaning that a cell habituated to light pulses will still be fully responsive to mechanical pulses, and vice versa. We therefore induced cells to contract by means of a light pulse (Figure 3F), turned off the light pulse to allow cells to re-extend for a fixed period of time, and then applied a mechanical pulse (see Methods for details of this experiment). By varying the time interval between when the light pulse stops and the mechanical pulse is applied, and exploiting the natural variation in cell re-extension times (see Figure 3A), we obtain a dataset in which we can group cells into those that contracted upon mechanical stimulation and those that did not, where for each cell in each group we calculated the fractional re-extension of the cell (see Methods). As shown in Figure 3G, we see a clear difference between the cells that did or did not contract, such that cells that had re-extended by more than about 10% of their final length were already fully responsive.

For cells that showed less than 10% extension, it would be difficult to know whether they had or had not contracted just because they are already close to their contracted size, making the contraction movement much more subtle, and potentially missed by our visual inspection. So our data do not provide strong support for a refractory period at any degree of re-extension, but in any case clearly show that extension by more than 10% is already enough to allow cells to contract again. Given that this level of re-extension takes place in 30s or less (Figure 3A), the refractory period, if there is one, would have to be at most 30s which, according to our model, should produce a corner frequency of 0.03 Hz, which is an order of magnitude higher than what we observe in our measurements (Figure 2B).

We conclude that cell re-extension is unlikely to explain the observed corner frequency for two reasons. First, the type of response created by a re-extension based refractory period would lead to a roll-off of 20 dB/decade, which does not fit our measurements. Second, the refractory period of at most 30s during re-extension is too short to explain the corner frequency of 500 Hz.

### Can existing models for Stentor habituation explain the frequency response?

Given that the refractory period model does not seem to be able to explain either the corner frequency or the slope of the roll-off, we asked whether any other existing models for habituation in *Stentor*, when viewed in the frequency domain, might explain these features. As discussed above, four models have been presented for habituation in *Stentor* (Smart 2024; Gershman 2024; Eckert 2024; Rajan 2025), each of which can successfully account for not only the response decrement that defines habituation, but also the additional “hallmarks of habituation” including the qualitative observation that frequency affects habituation extent. Because all four models can apparently account for all the currently existing data, it has not been possible to discriminate among them. The measurement of the *Stentor* frequency response now provides a new dataset that any successful model must explain, potentially providing a new way to discriminate among existing or future models.

The first model we consider is that of Smart et al 2024, which is based on a low pass filter that produces an output function that is then used to determine whether or not to contract when a stimulus arrives (Figure 4A). The transfer function predicted from this model is derived in Methods and plotted in Figure 4B. Fitting this function provides a best fit with -19 dB/decade, very close to the value of -20 dB/decade we would expect from the fact that the model is based around a one-pole low pass filter. Adding additional filter stages can increase the slope to 40 dB/decade, but there is no way to achieve 30. We thus argue that this model does not fit the data, at least not in its current form.

**Figure 4.**
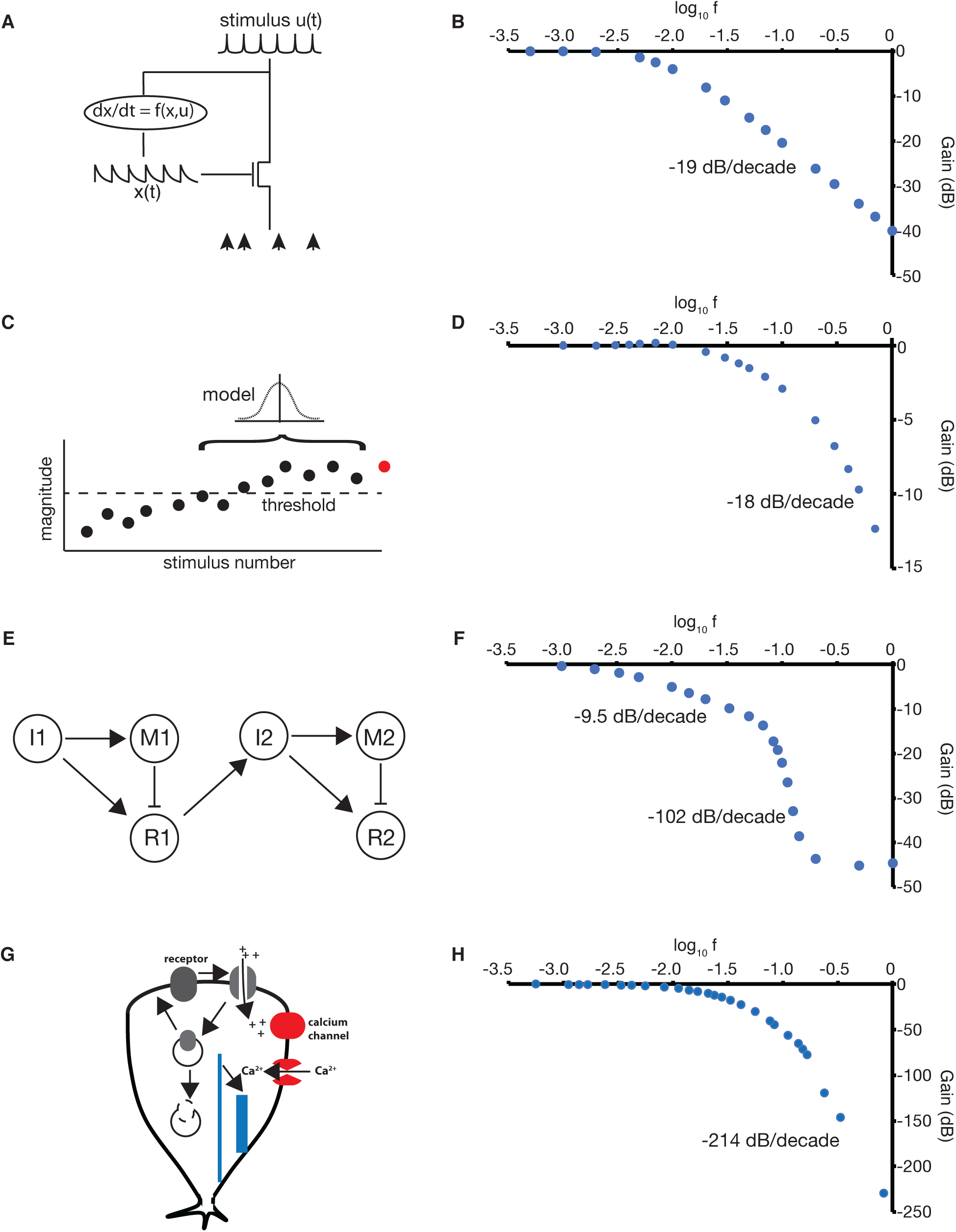
Transfer function predicted by existing models. **A**. convolution filter model (Smart 2024) in which the input u(t) is convolved with an impulse response (kernel) to produce a time-varying internal reference x(t), which is then used as a parameter in a function that determines whether or not the cell will respond to a stimulus. **B**. Frequency response predicted from convolution model using parameters for single stage filter given in Smart et al (2024) as described in Methods. **C**. Gaussian estimator model (Gershman 2024) in which the past history of stimulus data is used to update an internal model of the stimulus distribution, which is then used to determine the probability that a new stimulus exceeds a threshold of danger that would signal the need to contract. **D**. Frequency response of the Gaussian mixture model using software published in (Gershman 2024) as described in Methods. **E**. Concatenated Incoherent Feed Forward (CIFF) model in which two incoherent feed forward networks are chained together with the output of one driving the input of the other, as described in Eckert et al 2024. **F**. Frequency response of the CIFF model using parameters from Eckert et al (2024) as described in Methods. **G**. Receptor inactivation model in which it is proposed that mechanoreceptors that become stimulated can be inactivated via internalization and then either recycled back to the surface or else degraded, with active receptors creating an ion flow that changed membrane polarization and triggers voltage gated calcium channels that trigger contraction, as described by Rajan and Marshall (2025). **H**. Frequency response of the receptor inactivation model using parameters previously published (Rajan and Marshall 2025), as described in Methods. For each model, the best-fit slopes at high frequency are indicated on the plots.

The second model we consider is the Gaussian estimator model (Gershman 2024) (Figure 4C). We modified the algorithm in that paper as described in Methods to generate the transfer function across a range of frequencies, as plotted in Figure 4D. The shape of this curve is clearly different from the linear roll-off seen in our data. At higher frequencies the curve becomes more linear with a slope of -18 dB/decade, which is clearly different from the observed slope of 30. We conclude that the Gaussian estimator model, at least in the specific form published, does not account for the measured frequency response. We note that Gaussian estimators are just one of many possible estimators that one could design, and thus our data cannot rule out this general framework for explaining habituation.

The third model we consider is the concatenated incoherent feed-forward network model (Figure 4E). By modifying the program provided by the authors (Eckert 2024) to sweep a range of input frequencies, as described in Methods, we obtain the transfer function plotted in Figure 4F, which displays a roughly linear roll-off with a slope of - 10dB/decade over much of the frequency range, again inconsistent with the measured slope of -30.

Finally, we consider the model we have previously published that is based on receptor inactivation, internalization, and recycling (Figure 4G; Rajan 2025). As shown in Figure 4H, this model does not predict a sharp corner, and while it eventually reaches a linear roll-off, the slope is 214 dB/decade, much higher than what is seen in the observed transfer function. We conclude that this model also cannot explain the observe frequency response.

We had previously considered a simpler version of the receptor inactivation model in which receptors can be internalized and recycled back to the surface, but not degraded. In this case, the model does predict a clear corner and a linear roll-off, but now the roll-off is 20 dB per decade, consistent with the first-order kinetics with which internalized receptors are recycled back to the surface, which generates an exponential impulse response identical to that invoked in the Smart et al model. This is closer to 30 dB/decade but still not the same, and in any case we previously found that the model without degradation cannot account for many of the other habituation hallmarks (Rajan 2025).

We conclude that none of the existing models accounts for the 30 dB/decade linear roll-off, at least not in their current form. This result in itself does not prove that some other configuration or parameter set for a given type of model could not be found that would give the observed response. The Gaussian estimator model is just a specific instance of a very general class of estimator based models for habituation, and so the fact that this particular instance does not show the observed frequency domain does not rule out the possibility that some other form of estimator model might work. The concatenated incoherent feed forward model is a specific case of a general case of signaling network models which can have arbitrarily large numbers of nodes with various types of nonlinear functions connecting them. It is again thus not possible to rule out variations of the model that would explain our data, even if the current version does not appear to do so. But for the other two models presented here, namely the convolution kernel model (Smart 2024) and receptor inactivation (Rajan 2025) a formal argument shows that they are not able to produce a 30 dB/decade roll-off, no matter what parameters might be chosen. This is because both models are ultimately based on a system of one or more linear ordinary differential equations. It is a standard result in signal processing that filters built on linear ODEs have rational transfer functions, that is, transfer functions that can be expressed as the ratio of two polynomial functions of \ frequency. At sufficiently high frequency, the behavior of these polynomials is determined entirely by the term with the highest exponent, and the slope of the roll-off is set by the difference between the highest exponent in the numerator and denominator, which by definition is always an integer. For every difference of 1, the roll-off increases by 20 dB/decade, such that the slope will always be an integral multiple of 20. The 30 dB/decade slope that we observe is thus not realizable by any filter consisting of linear differential operators applied to the input and output, which is true for both models.

### A model for apparent fractional order frequency response

The 30 dB/decade roll-off that we observed is characteristic of a “fractional order” system, that is to say, one in which the transfer function is no longer a ratio of two polynomial functions of complex frequency with integer exponents. Fractional order behaviors typically arise in physical systems due to processes such as diffusion or percolation through fractal networks (Biswas 2017). In electrical systems, fractional order behavior can arise through similar processes as well as in “non-lumped” systems like transmission lines. We can potentially imagine various cellular mechanisms based on diffusion or transport through complex networks that could produce a fractional order response. These could potentially represent interesting new modes for intracellular signal processing that would contrast with standard biochemical reaction networks.

But might there be a way to implement the fractional order response using more conventional biochemical reactions? In analog circuit design, it is possible to emulate a fractional order roll-off by means of a “ladder” of filters arranged in parallel, whose corner frequencies (poles) span the frequency range over which the roll-off occurs, and whose gains decrease with increasing frequency in such a way that the combined effect produces a series of shoulders that follow a 10 dB/decade curve, with the overall effect of mimicking a true fractional order response with some ripples. Algorithms exist (Charef 1992; Freeborn 2010) to choose the poles and gains so as to maintain a constant phase difference across the frequency range. In our case, we are only interested in the gain, and so we simply look for a molecular mechanism that could produce a similar parallel array of filters.

One way to do this (Figure 5A) is to have a collection of different molecular species, each of becomes switched into an activated state by the stimulus, and while in their activated state, they inhibit the overall response of the system. Each of these molecular species would undergo an exponential decay back to the inactive form according to a rate constant that would be different for the different molecular species. This is equivalent to the different RC combinations in the parallel filters of the ladder in the analog circuits mentioned above. While in the active state, each molecular species inhibits the system response, but with a different efficacy, which would act like a gain for the corresponding filter in the ladder of an electronic circuit. By having all of the molecules responding to the same inputs, and feeding into the same outputs, they are effectively arranged in parallel like in the electrical filter ladder.

**Figure 5.**
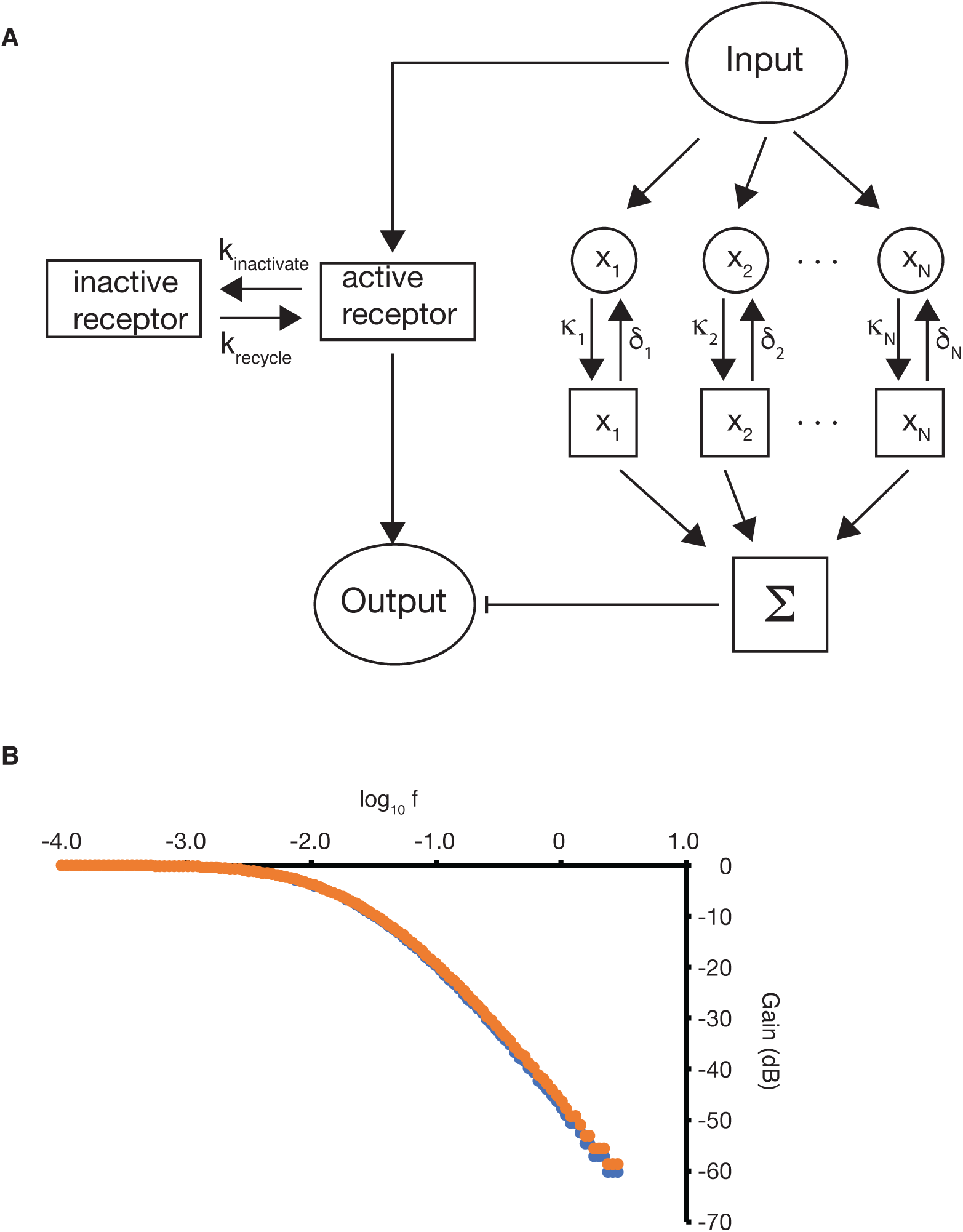
Model for fractional-order habituation response. **A**. Diagram of a hypothetical model in which a parallel set of inhibitory molecules act like the parallel filters of a ladder circuit to emulate a standard method for producing fractiona response in analog circuits. Each molecular species can be in either an inactive state denoted by a circle or an activated state denoted by a square. The outputs of these different species sum to produce an inhibitory inhibit that acts on the response pathway. **B.** Simulation of model with 12 species in the ladder with gains and rate constants forming a geometric series. (orange) transfer function with rate constants chosen as detailed in Methods (slope = 29.6 dB/decade). (blue) transfer function with rate constants manually optimized (slope = 30.3 dB/decade)

Intuitively, the way this model works is that whenever a stimulus arrives, all of the different molecules are stimulated into their active state, in which state they inhibit the system from responding to a second stimulus. But then over time, each molecular species population decays back to its inactive form. Those with the largest rate constant for decay will become inactive most quickly, while those with slower decay rate constants will stay active for longer. By the time a second stimulus arrives, some of the molecular species will have all returned to their inactive form, while some will remain active. The sooner the next stimulus arrives, the more of these inhibitory molecules will still be in their active form, and thus the more the output will be inhibited. The result is a low pass filter. Each of the individual inhibitory molecules is acting like a one-pole low pass filter, but with different corner frequencies and different gains. If we choose these corner frequencies and gains correctly, the parallel combination of filters can emulate a fractional order low pass filter.

Our previous experimental studies have provided evidence that calmodulin-dependent protein kinase II (CaMKII) plays a role in the habituation process. CaMKII is famous not only for its conserved role in learning in animal nervous systems, but also for the fact that it can exist in a large number of different phosphorylated isoforms generated through complex activity-dependent autophosphorylation processes (Hudmon 2002; Bayer 2025). The CaMKII complex exists as a dodecamer, which can either consist of a single type of subunit (homododecamer) or distinct types (heterododecamer) and has the ability to swap out monomers and exchange them between complexes in response to activation (Bhattacharyya 2020). We therefore hypothesize that in our system, CaMKII can be phosphorylated in response to a stimulus into any of 12 distinct phosphorylation states, each of which then has its own unique rate constant for returning to the dephosphorylated state, and each of which inhibits the contractile response of the cell with a different effective “gain”. We therefore implement a version of the model in Figure 5A in which there are twelve different molecular species, each corresponding to a different modified form of CaMKII.

In the particular version of the model presented here, the mechanoreceptors on the cell surface respond with some probability to a stimulus, and in turn send signals both to a downstream target such as a voltage gated calcium channel that triggers the contraction, as well as to CaMKII to promote its phosphorylation, with CaMKII then sending its inhibitory signal to the downstream target of the receptor. In this model, the CaMKII phosphorylated forms depicted in the right branch of Figure 5A can be viewed as forming an incoherent feed-forward loop, a point we will return to in the Discussion below. As with our previous biochemical model for habituation, we also make the assumption that the receptor itself can become auto-inactivated upon stimulation, for example by internalization into the cell where it no longer responds to stimuli, and can also be recycled back to the surface, as indicated on the left branch of Figure 5A.

With these assumptions, we simulated the steady state habituated response of the system as a function of frequency (see Methods for details). The result, as shown in Figure 5B, is a low pass filter with a flat passband, a single corner, and a linear roll-off with slope 30.3 when rate constants were manually tuned, and 29.6 dB/decade when rate constants were chosen according to the equation derived in Methods. In both cases, the decay times and the gains of the individual phosphorylation forms followed a geometric series, such that forms with a higher corner frequency had a lower gain, with the geometric constants chosen to achieve 30 dB/decade. This establishes that the simple scheme showed in Figure 5A can, at least in theory, be sufficient to produce the fractional order response that we have observed.

The key to obtaining a fractional order roll-off is that for each of the molecular species acting in parallel (in this case, CaMKII phosphorylated forms), the rate at which they return back to the unmodified form (i.e. become dephosphorylated) and the gain with which they inhibit the downstream target must be mathematically related such that as the corner frequency (decay rate) increases, the gain decreases proportionally. This may seem like an unrealistic assumption. However, if we assume that the phosphorylated form has a stronger effect on its targets the longer it remains in the phosphorylated state, then we see that doubling the decay rate, which means doubling the corner frequency, would directly cause the lifetime of the active state, and thus the effective gain, to be reduced by a factor of two.

We emphasize that the model presented here is likely just one of many different possible ways that a fractional order response could be realized. Our goal is not to identify the unique mechanism that is at work, but just to demonstrate that it is possible to obtain such a response from a biochemical network with a level of complexity that is commensurate with some of the known players in *Stentor* habituation like CaMKII. This is meant only as a starting point for further investigation.

## Discussion

### Molecular implementations

In introducing our hypothetical model to explain the fractional order response of the *Stentor* cell, we suggested the different molecular species in the “ladder” part of the circuit represent different phosphorylations of the CaMKII complex. This would be consistent with the known highly complex phosphorylation states that can occur in this large dodecameric protein complex.

But one could easily devise many alternative implementations, such as: multiple different receptors that can each be modified with some reversible post-translation modifications; distinct post-translational modifications on the same receptor that each inhibit its response and are reversed at different rates; multiple different kinases and/or that can each modify the receptor or some other key component, etc. What all of these types of mechanisms would have in common is that there are a set of biochemical changes that can happen independently of each other, each of which contributes to an overall inhibitory effect on the response with different efficacies, all of which are triggered by the stimulus (or some downstream signal), and which decay back to an inactive form with different time constants. Among all of these possibilities, we have only presented one hypothetical model and do not in any way make the claim that this is the only way the system could work.

Moreover, the whole scheme of a molecular “ladder” circuit of parallel molecular species is just one possible type of circuit topology that could create a fractional response. For example, we note in Figure 4D that the frequency response of the concatenated incoherent feed forward loop model also shows a fractional order, in this case a roll-off close to 10 dB/decade, suggesting that such networks have the potential to generate fractional order responses. It would be interesting to apply network enumeration methods (e.g. Ma 2009) to look for topologies that can create a range of fractional order responses.

### What sets the corner frequency?

With respect to the corner frequency, we do not currently know what process determines the relevant timescale. Our data argues that it is unlikely to be the extension of the cell following a contraction, but many other possibilities remain. As just a few examples, we can imagine that a time scale of a few hundred seconds could be set by synthesis of a protein, transport of a molecule through the cytoplasm of the giant cell, or some form of cytoskeletal remodeling independent of cell extension. In the model presented in Figure 5, the timescale/corner frequency is set by the slowest transition rate among the parallel molecular species that regulate the final output.

### Hidden variables in habituation

In this work we are invoking concepts from signal processing and describing the cell in terms of the transfer function of a filter. But it is important to note that the measured transfer function relates the frequency at which a series of pulsatile inputs are delivered to the probability of responding to each impulse, once the system has reached a steady state habituated response. Because both the input and outputs are in the form of a sequence of inputs, the overall system is not simply a linear filter, otherwise the output would be a superposition of impulse response functions produced by the filter. The actual filter links the mechanical input to some internal hidden variable, which would be calcium concentration, phosphorylation level of some protein, etc., and it is that hidden variable which in turn determines whether or not the cell will contract when the next stimulus arrives. A logical next step towards dissecting the mechanism of habituation will be to identify the internal hidden variables that underly the process, and then carry out a similar frequency domain analysis of how their levels or activities are linked to inputs of varying frequency. Given that fact that *Stentor* contraction is driven by calcium acting on the centrin cytoskeleton, the observation of calcium channels acting during the response, the effects of changing extracellular calcium on habituation (Rajan 2026), and the fact that the two molecules identified as playing a role in habituation (CaMKII and a novel EF-hand protein; Rajan 2026) are both putative calcium binding proteins, a major candidate for the internal hidden variable is clearly the intracellular calcium concentration. Other likely candidates include CaMKII activity and/or phosphorylation state, and any changes to the localization or post-translational modifications of the receptor itself. Once methods to measure these variables are available, the question will be to ask how their value at steady state habituation relates to the frequency of the stimuli.

### Functions of fractional order systems

Fractional order frequency responses have been observed in many different physical systems, and have been deliberately implemented using specialized electronic devices as well as emulated by networks of more conventional analog circuit elements (reviewed in Biswas 2017). In an engineering context, fractional order responses have been shown to confer improved performance of feedback control systems. Such systems can be more stable then integer order equivalents (Radwan 2009; Gulgowski 2026), and can track a control signal more quickly and with less overshoot (Bohannan 2008).

Within the general context of biological control systems, it will be interesting to explore whether fractional order behaviors may confer similar advantageous behaviors. Much of the literature on fractional order responses in biology pertains to the tissue and organ scale, with a focus on viscoelastic mechanics and distributed electrical activity (Magin 2004; Doehring 2005; Ugarte 2024).

Strikingly, fractional order frequency responses have been measured for sensory adaptation in retinal cells (Thorson 1974) as well as neural adaptation, measured both in single neurons (Lundstrom 2008; Lundstrom 2010) and in brain EEG signals (Lundstrom 2023). Given the fact that adaptation and habituation are fundamentally similar processes, differing only in where and how they are deployed and possibly not sharing all of the same additional hallmarks, the fact that we also see a fractional order response for the single-cell habituation of *Stentor* suggests this may be a general theme in adaptive/habituating systems at the single-cell level.

### Relevance of frequency domain properties for survival in the wild

Here we have mainly used the frequency domain analysis as a way to probe potential mechanisms of habituation, but this same information may also be telling us something about the biological role of habituation. Stimuli arriving a frequency lower than the corner frequency are essentially unable to cause habituation, meaning that such infrequent stimuli would always be viewed as something novel or threatening, such that the cell will always respond. One possible interpretation is that when stimuli are frequent, the cell has enough information about its environment to decide if it is safe or not, whereas when the stimuli arrive infrequently, the cell is not able to make a strong prediction about the safety of the environment and so the best strategy might to be contract the next time a stimulus appears. It would be interesting to know more about how the actual frequency of mechanical impacts experienced by a *Stentor* cell living in the pond might depend on the density of predatory organisms like rotifers or non-predatory prey organisms like swimming algae. We are unaware of such studies for *Stentor* or other protists but feel it could be a productive direction for future research.

## Methods

### Cell culture

*Stentor coeruleus* cells were purchased from VWR (Ward’s Science 470176-586; Radnor, PA) and grown in glass dishes in pasteurized spring water (PSW; Carolina Biological 132458; Burlington, NC). Cultures were fed every three days with *Chlamydomonas*, and detritus removed from the cultures daily. For all experiments, cells were washed three times with PSW and placed in a 35 mm polystyrene petri dish (#L331/10Benz Microscope Inc., Manchester MI) coated with poly-ornithine (Sigma) and then allowed to adhere stably for two days, with feeding with *Chlamydomonas* taking place on the second day. On the third day, the experiments (frequency response or elongation rate) were performed as described below.

### Measuring frequency response

The frequency response of habituation was measured using the stepper-motor based habituation device previously described (Rajan 2023b). In this device, the dish is placed on a metal ruler with 2 cm between the edge of the dish and the end of the ruler. Prior to each experiment, the cells on the dish were allowed to rest undisturbed for at least 4 hours, a timescale that prior experiments have shown is substantially longer than the time required for a cell to recover full response after being habituated by any stimuli. To acquire a response probability for any given frequency, the habituation device software was set to deliver pulses at a time interval equal to the reciprocal of the frequency. Images were acquired using a USB microscope (Celestron 44308; Torrance, CA), with image acquisition starting before turning on the habituation device in order to capture the initial response of the rested cells, which we take as the zero frequency response of the filter. Habituation was then carried out until the cells reached a steady state response.

Based on prior experiments, steady state was achieved within 30 min with taps delivered once per minute, and more rapidly with higher frequency taps. In each case, once steady state was achieved after a given number n of taps, a further n taps were then applied to ensure that the cells had reached a steady state response.

In order to calculate the response probability, video frames from the last 10-20 taps were analyzed visually and at each time point, the number of cells that were elongated prior to the tap, and the number of those cells that had contracted during the tap, were recorded, such that we scored the fraction of elongated cells that contracted during the tap. This fraction was then averaged over the last ten taps in the sequence. For frequencies higher than 0.5 Hz, the response probability is too low to observe more than a few contractions during any sequence of 10 stimuli. Therefore, we scored the last 30 taps for frequencies in the range 0.03-0.1 Hz, and the last 100 for frequencies above 0.1 Hz. Even using this procedure, the fraction of cells contracting at the highest frequency (1 Hz) was so low that the statistics become unreliable, hence although we show those datapoints on our plot, we did not include that frequency in fitting the slope of the roll-off.

### Measuring kinetics of cell re-elongation

To measure elongation after mechanical stimulation, the dish was placed on the stage of a Zeiss Stemi-2000 and video images acquired using a Celestron-HD 5MPI Eyepiece camera. After acquiring pre-tap images of the dish, the dish was briefly shaken, causing cells to contract. Video images were continuously acquired and lengths measured at time points using Fiji software. Distances were calibrated using images of a ruler acquired on the same day as the video.

### Testing for refractory period during cell re-elongation

Cells were induced to contract using light stimulation to avoid habituating effects on the mechanical response. This was accomplished by shining light from an LED flashlight (Baibian XHP160.6) located 3 cm from the surface of the dish for 20 seconds. The LED produced a light intensity of intensity 690 lx at a distance of 30 cm (measured with Sekonic C-4000 light meter), from which we calculate that cells were exposed to a light intensity of 7000 lx. The majority of cells contracted at the moment the light was removed. Cells were then allowed to re-extend for time intervals in the range 10-40 seconds after which a mechanical tap was applied using the standard habituation device with a force level of 5 (low force). Only cells that were extended prior to the light pulse and contracted after the light pulse were analyzed. Cell length was measured using Fiji before the light pulse (L_before_), after the pulse (L_after_), and before the mechanical stimulus (L_premechanical_), and the fractional elongation calculated according to (L_premechanical_ - L_after_)/(L_before_ - L_after_). Cells were visually scored for contraction following the mechanical stimulus and classified into either non-contracting or contracted. Values for fractional elongation were then plotted for cells in each of the two categories.

### Deriving frequency response for refractory period model

We consider the simplest model in which an isolated stimulus will cause contraction in 100% of cells, but if a second stimulus arrives during some fixed period after a cell has contracted, the cell will not contract. We denote this refractory period as 1_refractory_. We consider the typical experiment in which stimuli arrive periodically at a frequency f.

Intuitively we can see that when f is smaller than 1/1_refractory_, the refractory period will have no effect on the response, since it will always have elapsed before the next stimulus arrives. Once f exceeds 1/1_refractory_, then some stimuli will be arriving within 1_refractory_ after a preceding stimulus, and will thus be blocked from contracting. Since the time between successive stimuli is 1/f, for sufficiently high frequency the number of “blocked” stimuli during a given refractory period is given by

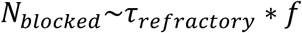

Since each refractory period was triggered after one stimulus that caused a contraction, the overall fraction of stimuli that cause a contraction is

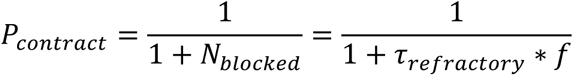

Viewing P_contract_ as the output, and f as the frequency of the input, we can see that this equation is identical to the amplitude of the transfer function of a one-pole low pass filter with a slope of 20 dB/decade and a corner frequency of 1/1_refractory_.

To simulate the refractory period model, we started with a simple simulation in which the probability of responding without the refractory period was 0.9 and the probability of responding during the refractory period was 0. This simulation is plotted in Figure 3D. We then extended the simulation to include a non-zero probability of contracting during the refractory period, with results plotted in Figure 3E. These simulations work by starting a counter any time a cell contracts and using a different response probability until the counter times out, at which point the response probability is switched back to the non-refractory probability.

### Derivation of frequency response for previously published habituation models

#### Gaussian Estimator Model

In order to obtain the steady state response as a function of frequency for the Gaussian Estimator model (Gershman 2024), the program accompanying the paper, habituation.ipynb, was modified to calculate the response at the range of inter-stimulus intervals spanning from 0.001 seconds to 1 second. The number of iterations of the simulation was increased to 100 to ensure that each run had the chance to reach its steady state response, which was then stored and output as a fraction rather than a percent. In order to have the program generate data in terms of actual time units, we scale time such that a length scale of 1 in the program corresponds to 100 seconds. The program was run in jupyter notebooks using parameters length _scale = 1, alpha = 0.3, threshold = 0.5.

#### Convolution Kernel Model

The convolution model of (Smart 2024) is based on an input stimulus sequence v(t) driving variation of an internal variable x(t) whose dynamics are given by

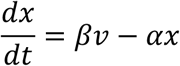

This internal variable determines the response y of the system to the current input stimulus according to

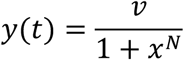

We can see that x can be obtained by convolving the input v by the impulse response

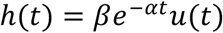

Where u is the Heaviside step function. For a standard habituation experiment, we assume the input stimulus sequence is a unit Dirac comb with spacing T=1/f. When the ith stimulus arrives, x was initially at a value xi, and then this is augmented by the incremental value β, and then this value undergoes an exponential decay to produce the value of x at the i+1th stimulus, hence,

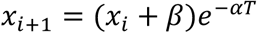

Once the system has reached steady state,

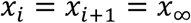

Solving for x_∞_ we obtain

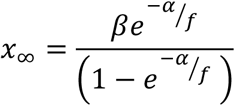

This is equivalent to equation 9 of (Smart 2024) which was derived by assuming an input sequence of rectangular pulses which in the limit of zero duty cycle becomes a Dirac comb. To generate our figure, we calculate x_∞_ for a range of frequencies, and then calculate the response y assuming an input comb has an amplitude of 10, β=1 and α=0.01 (as per Smart et al. 2024).

#### CIFF Model

In order to generate the frequency response at steady state for the concatenated incoherent feed forward loop model (Eckert 2024), the code used in the paper (Paper_figures_all.ipynb) was modified such that in addition to computing the habituation time, the response magnitude to the first and last stimuli were calculated within module plot_freq_or_is and used to compute the normalized steady state response. The parameter set used corresponded to that shown in Figure 3 of (Eckert 2024). Stimulus amplitude A was set to 10, and the inter-stimulus time T was varied between 1 and 1000. These are all in arbitrary units hence were plotted as such in Figure 4F.

#### Receptor Inactivation Model

In order to generate the frequency response at steady state for the receptor inactivation model (Rajan 2025), the same simulation program and parameters were used as in the 2025 paper but with a stimulus magnitude of 1 and inter-stimulus times varying from 0.01 to 30 minutes.

### Model for fractional order response

In order to implement a model for fractional order response based on a parallel set of molecular species, we assume a fixed number of receptors with a fixed probability of a given receptor being activated by the stimulus. When activated, each receptor has a probability of switching to an inactive state, after which inactive receptors can become active again according to a recycling rate constant. This part of the model is similar to our previous receptor inactivation model (Rajan 2025) except that we ignore the complexities of the activation function and simply assume a constant probability per receptor of activation. In addition to the receptor itself, we assume N distinct molecular species, in this particular case N=12, each of which exists as a population within which individual instances of the molecule can switch between an inactive and an activated state. The switch occurs with a constant probability whenever a stimulus is applied.

Each of these molecules when in the active state can return to the inactive state according to a decay constant d_i_ that is different for the 12 different species. While in the active state, each produces an output proportional to the product of the number of activated molecules of each species and a gain constant G_i_ that depends on the species. These outputs are summed to produce the overall output S. The number of activated receptors is then multiplied by 1 - S/Smax where Smax is the summed output from the ladder that would be obtained if all molecules of all species were activated.

The key parameters for this model are the decay constants and gain for the 12 molecular species acting in parallel. Each species acting individually behaves like a one pole filter with a corner frequency set by the decay constant and a DC gain set by G_i_. We thus represent the overall ladder stage as n single pole filters arranged in parallel, each responding to the same input and with their outputs summing. Each filter obeys the equation

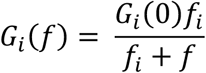

Where f_i_ is the corner frequency and G_i_ the gain for the ith filter. Following the approach taken in electronic fractional order circuits implemented with a ladder of filter stages (Charef 1992), we arrange the filters in a geometric series with frequency and gain scale factors α and β such that

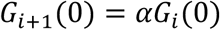

And

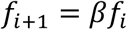

In order to obtain a design rule for choosing the gain and frequency scale factors, we take the usual linear approximation to a Bode function and assume that the corner frequency is where the gain just starts to drop below the zero frequency gain. Starting at the corner frequency f_1_ of the first filter, we can calculate the desired gain at the corner frequency of the second filter to produce a 10 dB/decade roll-off between the two poles:

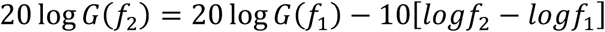

From which we obtain

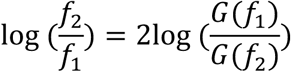

or

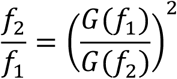

We make the approximation that since the first filter gain rolls off rapidly compared to the 10dB/decade we are trying to achieve, then by the time the second filter corner frequency is reached, the response is determined mainly by the second filter’s zero frequency gain such that G(f_2_) ∼ G_2_(0). Similarly, we assume that the zero frequency gain for the first filter, being larger than the second filter, dominates at lower frequencies such that G(f_1_) ∼ G_1_(0). This gives us:

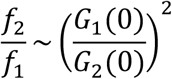

Substituting in the scale factors, we obtain:

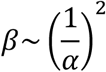

In practice, we first choose a value for beta that spreads the poles out over the desired bandwidth for the roll-off region up to the maximum frequency considered, and then choose alpha according to

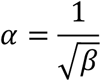

In the simulation plotted in Figure 5, β was chosen as 1.65 from which we obtain α = 0.78. The specific parameter set used for this plot was simulate_frequency_response_multiple_parallel_v7(1,100000, 0.1, 0.0001, 1.09, 120, 1.0, 0.1, 0.002, 0.0, 0.005, 0.1, 1.0, 1.65, 1.65, 1.0, 0.78, 1). Figure 5B also includes a second simulation in which the parameter α was manually tuned in attempt to get closer to a slope of -30, in this case using the value α=0.74. It can be seen that the rationally chosen parameters do essentially just as good a job as the manually adjusted ones.

## Acknowledgements

This work was initially supported by NSF grant MCB-2012647, then by NIH grant R35 GM130327, and John Templeton Foundation grant 63519. Work by students in the 2025 CCC Summer Course was supported by the Center for Cellular Construction, funded by NSF grant DBI-1548297. We acknowledge Deepa Rajan, Tatyana Makushok, Daniel Cortes, Sindy Tang, and Yogi Jaeger for enlightening discussions about learning in *Stentor*, Zhuoyi Song for a discussion of refractory periods in cell signaling, and William Bialek for pointing out the non-realizability of a 30 dB/decade response with linear lumped components.

## Author contributions

Built apparatus S.E., F.W., W.F.M.

Performed experiments S.E., A.M, F.W., E.M, A.B. K.R., G.K., J.M., K.B., W.F.M.

Analyzed data S.E., A.M, F.W., E.M, A.B. K.R., G.K., J.M., K.B., A.A., X.G.A. W.F.M.

Computational modeling W.F.M. Made figures W.F.M.

Wrote the paper W.F.M.

## Notes

### Competing Interest Statement

The authors have declared no competing interest.

## References

1. Baluška F, Levin M. (2016). On Having No Head: Cognition throughout Biological Systems. Frontiers in Psychology. 7, 902.

2. Bayer KU, Giese KP. 2025. A revised view of the role of CaMKII in learning and memory. Nat Neurosci. 28, 24–34

3. Bhattacharyya M, Karandur D, Kuriyan J. 2020. Structural Insights into the Regulation of Ca(2+)/Calmodulin-Dependent Protein Kinase II (CaMKII). Cold Spring Harb Perspect Biol. 12, a035147.

4. Binet A. 1897. The psychic life of micro-organisms. A study in experimental psychology. Open Court Publishing Co. Chicago. 120 pp.

5. Biswas K, Bohannan G, Caponetto R, Mendes Lopez A, Tenreiro Machado JA. 2017. Fractional-order devices. Springer Nature, Cham Switzerland.

6. Bohannan GW. 2008. Analog fractional order controller in temperature and motor control applications. J. Vib. Control 14, 1487–1498.

7. Bray D. (2011). Wetware: A computer inside every living cell. (Yale University Press, New Haven).

8. Charef A, Sun HH, Tsao YY, and Onaral B. (1992). Fractal system as represented by singularity function. IEEE Transactions on Automatic Control 37, 1465–1470.

9. Doehring TC, Freed AD, Carew EO, Vesely I. 2005. Fractional order viscoelasticity of the aortic valve cusp: an alternative to quasilinear viscoelasticity. J Biomech Eng. 127, 700–8

10. Dussutour A. (2021). Learning in single cell organisms. Biochemical and Biophysical Research Communications. 564, 92–102.

11. Eckert L, Vidal-Saez MS, Zhao Z, Garcia-Ojalvo J, Martinez-Corral R, Gunawardena J. 2024. Biochemically plausible models of habituation for single-cell learning. Curr. Biol. 34, 5646–5658.e3

12. Fernando CT, Liekens AML, Bingle LEH, Beck C, Lenser T, Stekel DJ, Rowe JE. (2009). Molecular circuits for associative learning in single-celled organisms. J. R.. Soc. Interface 6, 463–9

13. Ferrell JE. (2016). Perfect and near-perfect adaptation in cell signaling. Cell Systems 2, 62–67

14. Freeborn TJ, Maundy B, Elwakil A. (2010). Towards the realization of fractional step filters. Proceedings of 2010 IEEE International Symposium on Circuits and Systems, Paris, France, 2010, pp. 1037–1040, doi: 10.1109/ISCAS.2010.5537360.

15. Gershman SJ, Balbi PE, Gallistel CR, Gunawardena J. 2021. Reconsidering the evidence for learning in single cells. Elife. 2021 Jan 4;10:e61907. doi: 10.7554/eLife.61907.

16. Gershman SJ. 2024. Habituation as optimal filtering. iScience 27, 110523

17. Golding I. 2011. Decision making in living cells: lessons from a simple system. Annu Rev Biophys. 40, 63–80

18. Gulgowski J, Stefański TP. 2026. Stability of fractional-order systems using the Marchaud derivative with a response depending on an infinite time interval of history. Communications in Nonlinear Science and Numerical Simulation. 157,109787.

19. Gunawardena, J. (2022). Learning Outside the Brain: Integrating Cognitive Science and Systems Biology. Proc. IEEE 110, 590–612

20. Harris JD. (1943). Habituatory response decrement in the intact organism. Psychological Bulletin. 40, 385–422.

21. Hell JW. 2014. CaMKII: claiming center stage in postsynaptic function and organization. Neuron. 81, 249–65

22. Huang B, Pitelka DR. 1973. The contractile process in the ciliate, Stentor coeruleus. I. The role of microtubules and filaments. J. Cell Biol. 57, 704–28.

23. Hudmon A, Schulman H. 2002. Structure-function of the multifunctional Ca2+/calmodulin-dependent protein kinase II. Biochem J. 364, 593–611

24. Jennings HS. 1902. Studies on reactions to stimuli in unicellular organisms. IX. On the behavior of fixed infusoria (Stentor and Vorticella) with special reference to the modifiability of protozoan reactions. Am. J. Physiol. 8, 23–60

25. Jennings HS. 1906. Behavior of the Lower Organisms. Columbia University Press, New York, NY, USA.

26. Jones AR, Jahn TL, Fonseca JR. (1970). Contraction of protoplasm. III. Cinematographic analysis of the contraction of some heterotrichs. Journal of Cellular Physiology. 75, 1–7.

27. Kuenen LPS, Baker TC. (1981). Habituation versus sensory adaptation as the cause of reduced attraction following pulsed and constant sex pheromone pre-exposure. J. Insect Physiol. 27, 721–726.

28. Kukushkin, N.V., Carney, R.E., Tabassum, T., and Carew, T.J. (2024). The massed-spaced learning effect in non-neural human cells. Nature Communications. 15, 9635

29. Larson BT, Garbus J, Pollack J, Marshall WF. (2022). A unicellular walker controlled by a microtubule-based finite-state machine. Current Biology 32, 3745–3757.

30. Larson BT. 2023. Perspectives on Principles of Cellular Behavior from the Biophysics of Protists. Integr Comp Biol. 2023 Dec 29;63(6):1405–1421

31. Levin M, Keijzer F, Lyon P, Arendt D. 2021. Uncovering cognitive similarities and differences, conservation and innovation. Philos Trans R Soc Lond B Biol Sci. 376, 20200458.

32. Lisman J. 2017. Criteria for identifying the molecular basis of the engram (CaMKII, PKMzeta). Mol Brain. 10, 55

33. Lundstrom BN, Higgs MH, Spain WJ, Fairhall AL. 2008. Fractional differentiation by neocortical pyramidal neurons. Nat Neurosci. 11, 1335–1342

34. Lundstrom BN, Fairhall AL, Maravall M. 2010. Multiple timescale encoding of slowly varying whisker stimulus envelope in cortical and thalamic neurons in vivo. J Neurosci. 30, 5071–5077

35. Lundstrom BN, Richner TJ. 2023. Neural adaptation and fractional dynamics as a window to underlying neural excitability. PLoS Comput Biol. 19, e1010527

36. Lyon P. (2015). The cognitive cell: bacterial behavior reconsidered. Front Microbiol. 6, 264.

37. Lyon P, Keijzer F, Arendt D, Levin M. 2021. Reframing cognition: getting down to biological basics. Philos Trans R Soc Lond B Biol Sci. 376, 20190750

38. Ma W, Trusina A, El-Samad H, Lim WA, Tang C. 2009. Defining network topologies that can achieve biochemical adaptation. Cell. 138, 760–73

39. Magin RL. 2004. Fractional calculus in bioengineering. Crit Rev Biomed Eng. 32, 1–104

40. Maloney M, McDaniel W, Locknar S, Torlina H. (2005). Identification and localization of a protein immunologically related to caltractin (centrin) in the myonemes and membranelles of the heterotrich ciliate Stentor coeruleus. Journal of Eukaryotic Microbiology 52, 328–338.

41. Marshall WF. (2021). Regeneration in Stentor coeruleus. Front Cell Dev Biol 9, 753625

42. Nakagaki T, Yamada H, Toth A. (2000). Maze-solving by an amoeboid organism. Nature 407, 470–470.

43. Newman E. (1972). Contraction in Stentor coeruleus: A Cinematic Analysis. Science. 177, 447–449.

44. Odde DJ, Buettner HM. 1995. Time series characterization of simulated microtubule dynamics in the nerve growth cone. Ann Biomed Eng. 23, 268–86

45. Odde DJ, Buettner HM. 1998. Autocorrelation function and power spectrum of two-state random processes used in neurite guidance. Biophys J. 75, 1189–96

46. Radwan AG, Soliman AM, Elwakil AS. (2009). On the stability of linear system with fractional order elements. Chaos, Solitons, Fractals 40, 2317–2328.

47. Rajan D, Makushok T, Kalish A, Acuna L, Bonville A, Correa Almanza K, Garibay B, Tang E, Voss M, Lin A, Barlow K, Harrigan P, Slabodnick MM, Marshall WF. (2023a). Single-cell analysis of habituation in Stentor coeruleus. Current Biology. 33, 241–251.

48. Rajan D, Chudinov P, Marshall W. (2023b). Studying Habituation in Stentor coeruleus. J Vis Exp. 191, 10.3791/64692

49. Rajan D, Marshall WF. (2025). A receptor-inactivation model for single-celled habituation in Stentor coeruleus. Current Biology. 35, 3327–3340

50. Rajan DH, Albright A, Kim H, Diaz U, Hudnall Y, Steube N, Dey G, Liu T, Marshall WF. (2026). Molecular pathways for learning in the single-cell Stentor coeruleus. Curr Biol. 36, 2367–2381.e9

51. Rankin CH, Abrams T, Barry RJ, Bhatnagar S, Clayton DF, Colombo J, Coppola G, Geyer MA, Glanzman DL, Marsland S, McSweeney FK, Wilson DA, Wu CF, Thompson RF. (2009). Habituation revisited: an updated and revised description of the behavioral characteristics of habituation. Neurobiol Learn Mem. 92, 135–8.

52. Reid CR. 2023. Thoughts from the forest floor: a review of cognition in the slime mould Physarum polycephalum. Anim Cogn. 2023 Nov;26(6):1783–1797.

53. Ros-Rocher N, Brunet T. 2023. What is it like to be a choanoflagellate? Sensation, processing and behavior in the closest unicellular relatives of animals. Anim Cogn. 2023 Nov;26(6):1767–1782

54. Saigusa T, Tero A, Nakagaki T, Kuramoto Y. (2008). Amoebae anticipate periodic events. Phys. Rev. Lett. 100, 018101.

55. Silva HS, Martins ML, Vilela MJ, Jaeger R, Kachar B. 2006. 1/f ruffle oscillations in plasma membranes of amphibian epithelial cells under normal and inverted gravitational orientations. Phys Rev E Stat Nonlin Soft Matter Phys. 74, 041903.

56. Smart M, Shvartsman SY, Mönnigmann M. (2024). Minimal motifs for habituating systems. Proc. Natl. Acad. Sci. U.S.A. 121, e2409330121

57. Staddon JER, Higa JJ. (1996). Multiple time scales in simple habituation. Psychol. Rev. 103, 720–33

58. Tang SKY, Marshall WF. (2018). Cell Learning. Curr. Biol. 28, R1180–1184

59. Tartar V. (1961). The Biology of Stentor. Pergammon Press, NY.

60. Thompson RF, Spencer WA. (1966). Habituation: a model phenomenon for the study of neuronal substrates of behavior. Psych. Rev. 73, 16–43.

61. Thorson J, Biederman-Thorson M. 1974. Distributed relaxation processes in sensory adaptation. Science. 1974 183, 161–72

62. Torre V, Ashmore JF, Lamb TD, Menini A. (1995). Transduction and Adaptation in Sensory Receptors. J Neuroscience. 15, 7757–7768.

63. Trifonova K, Falk MJ, Rouches M, Vaikuntanathan S, Elowitz M, Murugan A. 2025. Trainable computation in molecular networks. bioRxiv [Preprint]. 2025 Dec 30:2025.12.28.696421

64. Uchida G, Chinzei T, Matsuura H. 1999. Reverse motion of organelles with myosin molecules along bundles of the actin filaments in a Characean internodal cell. Biochem Biophys Res Commun. 257, 223–7

65. Ueda T, Kobatake Y. 1983. Quantitative analysis of changes in cell shape of Amoeba proteus during locomotion and upon responses to salt stimuli. Exp Cell Res. 147, 466–71.

66. Ugarte JP, Tobón C. 2024. Fractional-order modeling of myocardium structure effects on atrial fibrillation electrograms. Math Biosci. 378, 109331

67. Wan KY, Jékely G. 2021. Origins of eukaryotic excitability. Philos Trans R Soc Lond B Biol Sci. 376, 20190758.

68. Wan KY. 2024. Biophysics of protist behaviour. Curr Biol. 2024 Oct 21;34(20):R981–R986

69. Wood DC. (1970a). Parametric studies of the response decrement produced by mechanical stimuli in the protozoan, Stentor coeruleus. Journal of Neurobiology. 1, 345–360.

70. Wood DC. (1970b). Electrophysiological studies of the protozoan, Stentor coeruleus. J. Neurobiol. 1, 363–377.

71. Wood DC. (1977). A curariform binding site mediates mechanical stimulus transduction in Stentor coeruleus. Comp. Biochem. Physiol. 56C, 155–166.

72. Wood DC. (1982). Membrane permeabilities determining resting, action and mechanoreceptor potentials in Stentor coeruleus. Journal of Comparative Physiology. 146, 537–550.

73. Wood DC. (1985). The mechanism of tubocurarine action on mechanoreceptor channels in the protozoan stentor coeruleus. J. Exp. Biol. 117, 215–235

74. Wood DC. (1973). Stimulus specific habituation in a protozoan. Physiol. Behavior 11, 349–354.

75. Vertosick FT. (2002). The Genius Within: Discovering the Intelligence of Every Living Thing. Harcourt, San Diego.

